# Recombinant HcGAPDH Protein Expressed on Probiotic *Bacillus subtilis* Spores Protects Sheep from *Haemonchus contortus* Infection by Inducing both Humoral and Cell-mediated Responses

**DOI:** 10.1101/872291

**Authors:** Yi Yang, Guiheng Zhang, Jie Wu, Xueqiu Chen, Danni Tong, Yimin Yang, Hengzhi Shi, Chaoqun Yao, Lenan Zhuang, Jianbin Wang, Aifang Du

## Abstract

Probiotic Bacillales have been shown effective in controlling pathogens. In particular, live probiotic bacteria are thought to improve the composition of gastrointestinal microbiota, and to reduce pathogen colonization. However, how probiotics regulate immune responses and protect the host from parasitic infection remains largely unknown. In this study, we investigated whether Bacillales can be used against *Haemonchus contortus*, a parasitic nematode that infects small ruminants in sheep and goats worldwide. Using 16S ribosomal RNA sequencing, we found that Bacillales was highly depleted in the abomasal microbiota of sheep infected with *H. contortus*. We constructed a recombinant *Bacillus subtilis* strain (rBS*^CotB-HcG^*) that express glyceraldehyde-3-phosphate dehydrogenase of *H. contortus* (HcGAPDH) on its spore surface. However, mice orally administrated with the rBS*^CotB-HcG^* strain showed strong Th1-dominated immune responses; and sheep administrated *per os* with rBS*^CotB-HcG^* showed increased proliferation of peripheral blood mononuclear cells, elevated anti-HcGAPDH IgG levels in sera, and higher anti-HcGAPDH sIgA levels in intestinal mucus. In addition, treatment of *H. contortus* infected sheep with rBS*^CotB-HcG^* (Hc+rBS*^CotB-HcG^*) promoted the abundance of probiotic species in the abomasal microbiota; it also improved the average weight gain of the sheep by 27.7%. These Hc+rBS*^CotB-HcG^* sheep have reduced number of eggs per gram of feces (by 84.1%) and worm burdens (by 71.5%), with alleviated abomasal damage by *H. contortus*. Collectively, our data demonstrate the protective roles of CotB-HcGAPDH-expressing *B. subtilis* spores against *H. contortus* infection, suggesting a potential value of using this probiotic-based strategy in controlling parasitic nematodes of socioeconomic importance.

**Importance:** Sequencing of the infected sheep ’s stomach flora revealed potential probiotics that could control *H. contortus* infection, and further genetically engineered recombinant probiotic spores expressing parasite protein, and validated their good immunogenicity in a mouse model. In the sheep infection model, the recombinant probiotics have proven to be effective against parasite infections.

## Introduction

*Haemonchus contortus* is one of the most economically important parasites causing haemonchosis in small ruminants around the world [1]. Haemonchosis may lead to anemia, weakness and even death of host animals prior to parasite’s pre-patent period [2, 3]. Anthelmintics have been the mainstay to control *H. contortus* infection. However, resistant *H. contortus* strains to widely used anthelmintics such as ivermectin are prevalent in many geographic regions [4]. Developing new prevention strategies against haemonchosis are challenging, although some progress has been made in addressing the mechanisms of *H. contortus* resistance [5]. Besides, residues of anthelmintics in meat, milk and other products impact human health and is a great public concern [6].

Probiotics are known to improve human and animal health. In particular, probiotics from food sources are thought to reduce intestinal infections by pathogens [7]. A previous study showed that *Bacillus subtilis* inhibited the colonization of *Staphylococcus aureus* by affecting its Agr quorum sensing system [7]. Bacterial spores can withstand extreme adverse environments with long-term survival rate. Therefore, the *Bacillus* spp. are considered as suitable probiotic candidates. In addition, *B. subtilis* spores have adjuvant effects [8, 9]. *B. subtilis* has recently been classified as the novel food probiotics for human and animal consumption and is widely used as oral vaccine vehicle [10], such as delivery of heterologous antigens to gastrointestinal tract as bioactive molecules [11, 12]. A recent study showed that CotC, a major component of the *B. subtilis* spore coat, was able to carry *Clonorchis sinensis* cysteine protease on the spore surface [13]. Recombinant *B. subtilis* spores expressing a tegumental protein was shown to provide protection against *C. sinensis* infection in a rat model [14].

HcGAPDH, an important component of *H. contortus* excretory/secretory products, is a glycolytic enzyme [15, 16]. In many organisms, GAPDHs are shown to have additional functions other than their enzymatic activity in glycolysis. It was shown that a recombinant HcGAPDH DNA vaccine can protect the recipient sheep from *H. contortus* infection by inducing effective host immune responses [17]. However, this DNA vaccine has not been applied in clinical practice, likely due to its limited commercial availability [18]. Given the prevalent anti-drug resistant for *H. contortus*, a better immune protection strategy against haemonchosis is still needed. The purposes of this study were to develop an oral vaccine using recombinant *B. subtilis* spores expressing CotB-HcGAPDH fusion protein and to investigate its underlying mechanisms. Overall, our data demonstrated a recombinant spore-based strategy as an alternative to anthelmintics.

## Results

### Relative abundance of Bacillales negatively correlated with *H. contortus* infection

To investigate the effect of microbiota on *H. contortus* infection, we analyzed abomasal microbiota of *H. contortus*-infected sheep using 16S ribosomal RNA (rRNA) gene sequencing. In the control sheep without *H. contortus* infection, the abomasal microbiota were dominated by the following bacterial class: Alteromonadales (35.5%), Pseudomonadales (29.5%), Bacteroidales (10.4%), Clostridiales (9.8%), Flavobacteriales (3.4%), Enterobacteriales (1.9%), Bacillales (1.3%), Aeromonadales (1.0%) (Fig 1a). *H. contortus* infection induced significant changes in microbial abundance including those of Alteromonadales, Pseudomonadales, Sphingobacteriales, Enterobacteriales, Bacillales, and Coriobacteriales, compared to the uninfected group (Fig 1a). Of particular interesting is the Bacillales group that has the probiotic effects in relation to *H. contortus* infection. We found that the relative abundance of Bacillales was significantly reduced after *H. contortus* infection (Fig 1b and 1c) (*p* < 0.005). In addition, our linear effect size (LEfSe) analysis on the 16S rRNA sequences showed that Bacillales is the main contributor as a probiotic in the abomasal microbiota to protect sheep from *H. contortus* infection (Fig 1d). Together, these data showed that sheep with *H. contortus* infection have led to significant reduction of Bacillales in the microbiota of abomasum, suggesting a potential protective role of these probiotic bacteria against pathogen infection.

**Fig 1.**
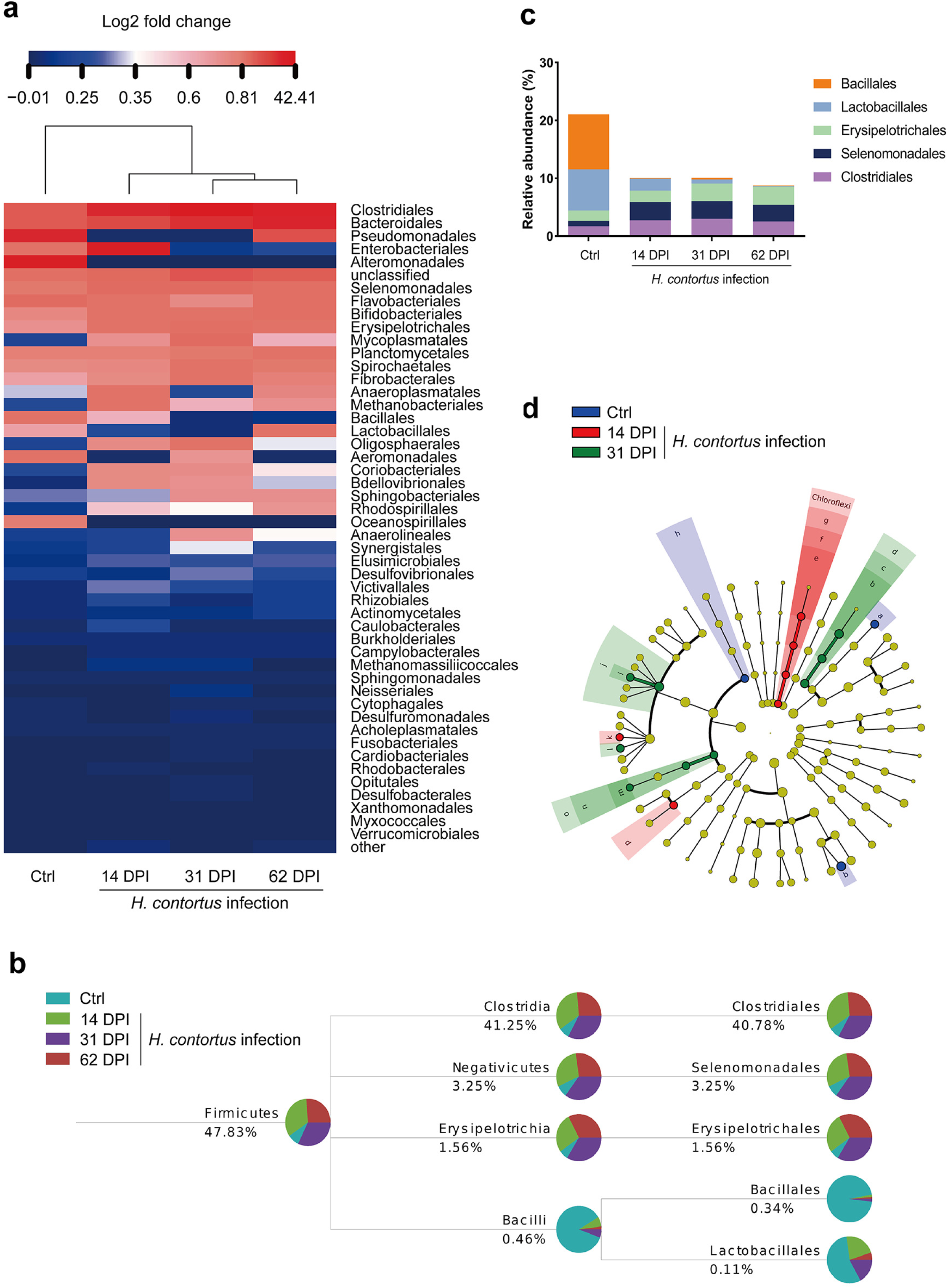
Relative abundance of Bacillales was related with *H. contortus* infection in sheep. **a.** Heatmap of relative abundance of abomasal bacteria. Color breaks in heatmap are adjusted to show relative abundance at <0.3% (blue shades), 0.3-0.4% (white shades), and >0.4% (red shades). DPI, Day post infection. **b.** Community taxonomic system composition analysis of Firmicutes. Relative abundance of abomasal bacteria in each sample is shown by a colored pie chart. **c.** The taxonomic composition of of Firmicutes. The proportion of different color blocks indicates relative abundance of different species. **d.** The taxonomic cladogram obtained by the linear effect size (LEfSe) analysis of 16S sequences within groups. Different colors represent different groups, and different color nodes in the branches represent groups of microorganisms that play an important role in the corresponding group of colors (a, Myroides; b, Sphingobacteriaceae; c, Sphingobacteriales; d, Sphingobacteriia; e, Anaerolineaceae; f, Anaerolineales; g, Anaerolineae; h, Bacilli; i, Lachnospiracea_incertae_sedis; j, Lachnospiraceae; k, Pseudoflavonifractor; l, Ruminococcus; m, Bulleidia; n, Erysipelotrichales; o, Erysipelotrichia; p, Veillonellaceae; q, Psychrobacter).

### Expression of CotB-HcGAPDH on the surface did not affect the production or the structure of *B. subtilis* spores

Based on the available data, we developed a protocol to generate recombinant spores expressing the fusion protein on the surface by joining the *B. subtilis* spore coat protein B (CotB) and the *H. contortus* GAPDH (HcGAPDH) protein (CotB-HcGAPDH or CotB-HcG) (Fig 2a). First, the full-length cDNA of *HcGAPDH* was cloned into the pET32a vector (pET32a-HcGAPDH), and the recombinant HcGAPDH protein was expressed and purified (Fig 2b and 2c). The purified protein was then used to generate polyclonal antibodies. Second, the *HcGAPDH* and *CotB* genes were fused and cloned into the pDG364 vector (pDG364-*CotB*-*HcGAPDH*). The fusion protein CotB-HcGAPDH (CotB-HcG) was expressed in *B. subtilis* spores (rBS*^CotB-HcG^*) (Fig 2d and Fig 2e).

**Fig 2.**
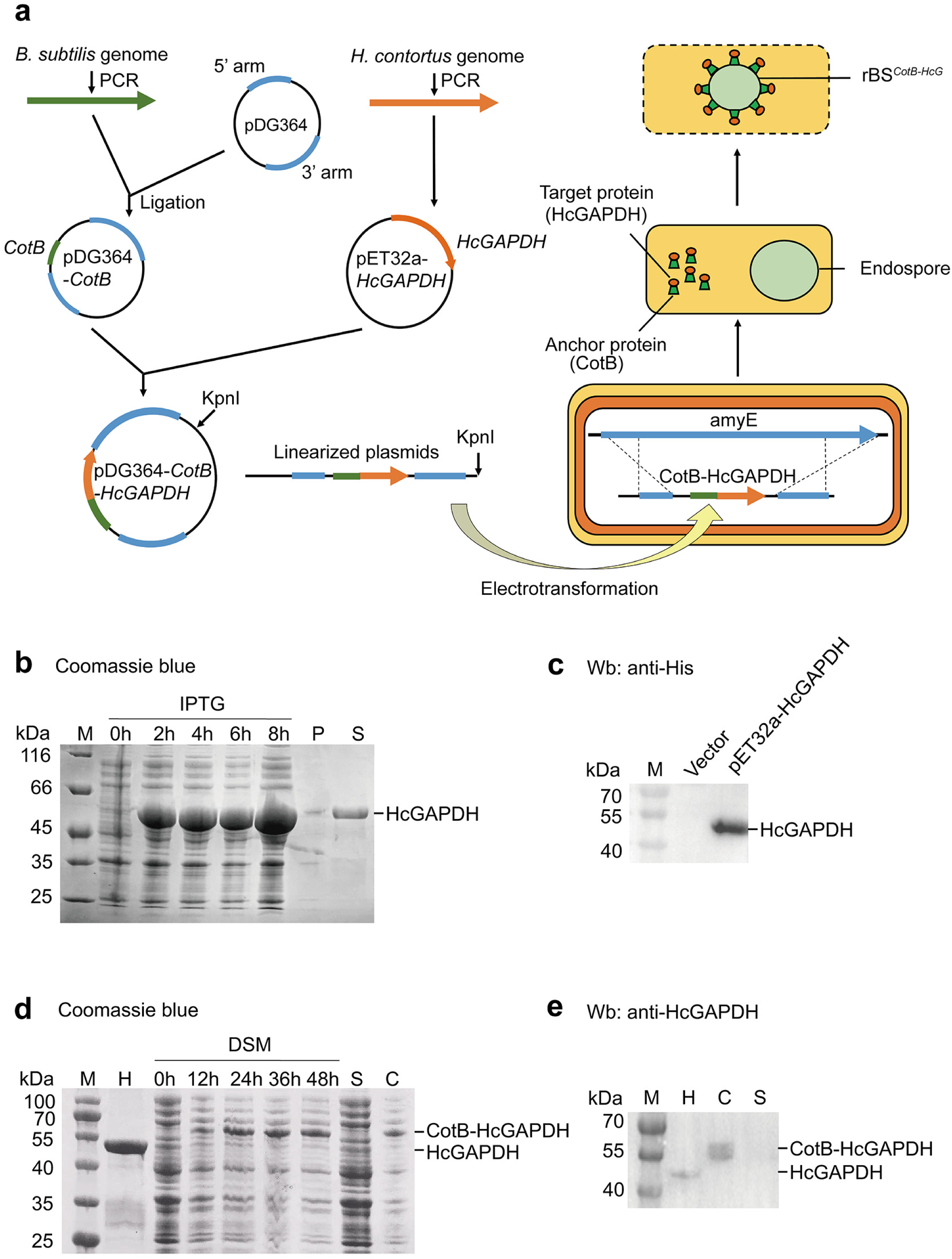
Recombinant *B. subtilis* spores expressing CotB-HcGAPDH on the surface. **a.** Schematic of genetic engineering to generate recombinant spores with CotB-HcGAPDH presenting on the surface (rBS*^CotB-HcG^*). **b.** Coomassie blue staining of the recombinant HcGAPDH protein. M, protein marker; HcGAPDH, recombinant GAPDH from *H. contortus*; P, pellet; S, supernatant; IPTG, Isopropyl β-D-Thiogalactoside. **c**. Western blotting of the recombinant HcGAPDH with anti-His antibody. Vector, empty pET32a control. **d.** Coomassie blue staining of the CotB-HcGAPDH fusion protein in *B. subtilis*. H, purified HcGAPDH protein; S, supernatant; C, spore coat from rBS*^CotB-HcG^; DSM,* Difco sporulation medium. **e**. Western blotting of the CotB-HcGAPDH fusion protein with anti-HcGAPDH rabbit antibody.

To verify that the recombinant fusion protein was expressed on the surface of *B. subtilis* spores, we have performed the immunofluorescence (IF) using the polyclonal antibodies to HcGAPDH on the bacterial spores induced in Difco sporulation medium (DSM). CotB-HcGAPDH in rBS*^CotB-HcG^* started to appear on the spore coat after 24 h of induction and increased steadily between 24 h and 72 h (Fig 3a). Flow cytometry (FCM) assay further confirmed that 86.01% of the rBS*^CotB-HcG^* spores expressed CotB-HcGAPDH 72 h after induction (Fig 3b). There was no difference in production and germination of spores between the wild-type and the rBS*^CotB-HcG^* strains (Fig 3c). To determine whether expression of CotB-HcGAPDH affected spore structure, rBS*^CotB-HcG^* spores were examined using scanning electron microscope (SEM) and transmission electron microscope (TEM). There was no change on coat folds of elliptical spore morphology between the wild-type and the rBS*^CotB-HcG^* strains (Fig 3d). In addition, TEM images revealed clear exine and intine structures of rBS*^CotB-HcG^* similar to the wild-type strain (Fig 3e). These results indicate that expression of CotB-HcGAPDH fusion protein did not change the production or the structure of *B. subtilis* spores.

**Fig 3.**
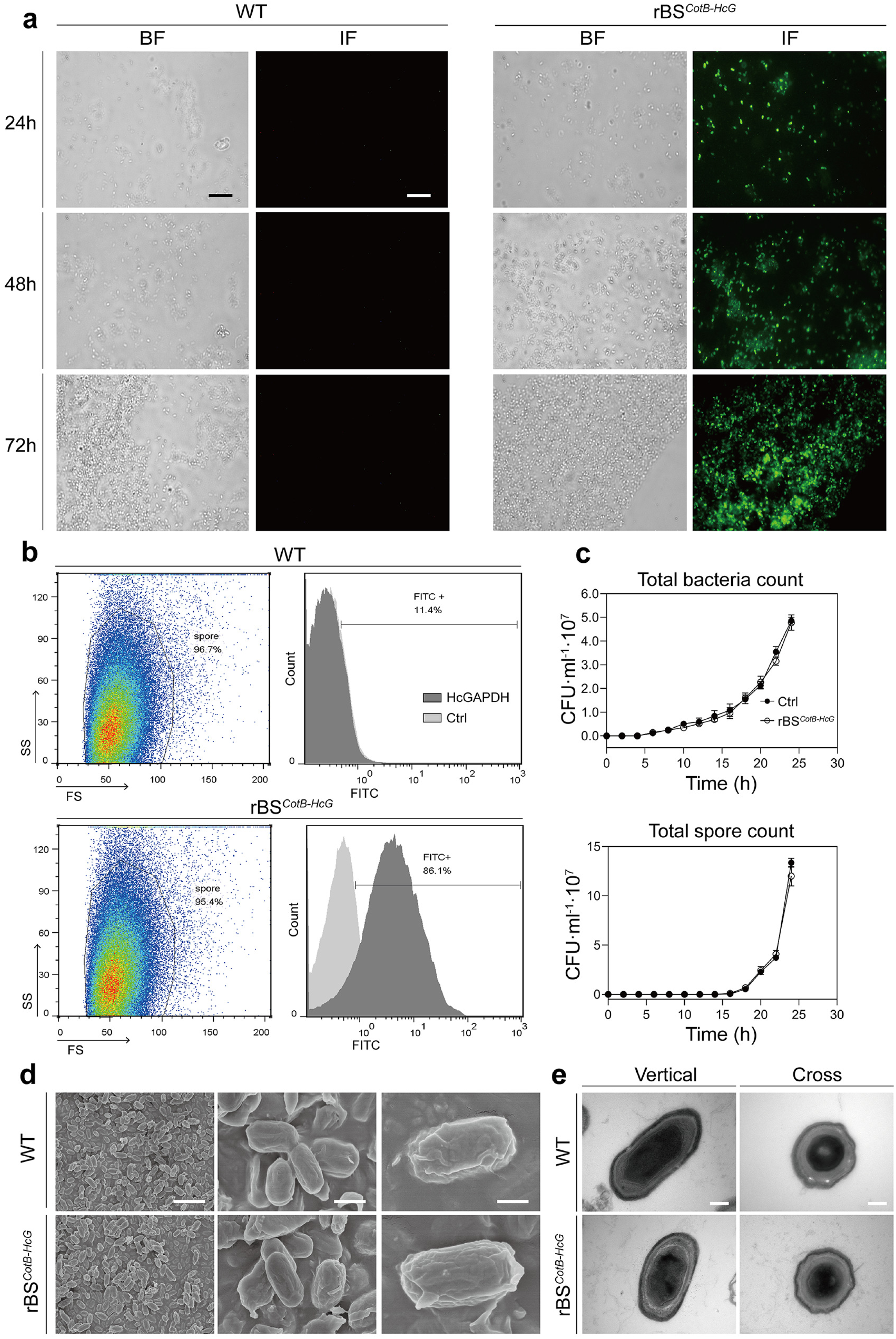
Expression of CotB-HcGAPDH fusion protein did not affect the structure and production of *B. subtilis* spores. **a.** Immunofluorescence (IF) of CotB-HcGAPDH expressed on the surface of wild-type (WT) and rBS*^CotB-HcG^* spores. 24 h, 48 h and 72 h indicate different time points after spore induction by DSM. BF, bright field; IF, immunofluorescence. Scale bar = 1 μm. **b**. Flow cytometry (FCM) analysis of CotB-HcGAPDH expression on the surface of WT and rBS*^CotB-HcG^* spores. FS, forward scatter; SS, side scatter. **c.** Production and germination analysis of WT and rBS*^CotB-HcG^* spores. **d.** Representative images of WT and rBS*^CotB-HcG^* spores by scanning electron microscope (SEM). Scale bar = 10 μm (left), 1 μm (middle and right). **e**. Representative images of WT and rBS*^CotB-HcG^* spores by transmission electron microscope (TEM). Scale bar = 200 nm (left), 100 nm (right).

### Recombinant *B. subtilis* spores expressing CotB-HcGAPDH fusion protein stimulated both humoral and cell-mediated immune responses in mice and sheep

To test whether the recombinant *B. subtilis* spores have positive probiotic effects in promoting immune responses, mice were orally administrated with PBS (Ctrl), wild-type (WT) strain, rBS*^CotB^* or rBS*^CotB-HcG^* spores, and purified HcGAPDH protein, respectively (Fig 4a). Lymphocytes prepared from the spleen samples from mice treated as above were cultured and stimulated with ConA, LPS or the purified HcGAPDH protein to examine the specific cell-mediated immune responses. Both ConA and LPS groups showed that rBS*^CotB-HcG^* administration showed higher levels of lymphocyte proliferation than the control group (*p* < 0.01) (Fig 4b). The purified HcGAPDH protein also stimulated lymphocyte proliferation with statistical significance as compared with the control group (*p* < 0.05). To investigate whether humoral immune responses were activated by the rBS*^CotB-HcG^* spores, we measured anti-HcGAPDH Immunoglobulin G (IgG) levels in the murine sera. We found that rBS*^CotB-HcG^* induced the highest antibody level (*p* < 0.005 as compared with Ctrl) at week 3 (Fig 4c). The purified HcGAPDH protein also induced higher level of specific antibody than the control group (*p* < 0.01). No anti-HcGAPDH antibody was detected in the mice receiving PBS, the wild-type or the rBS*^CotB^* strain (*p* > 0.05). The subtype IgG2a or IgG1 reflects whether the type of T cell immune response is dominated by Th1 or Th2, respectively [19]. To further determine the Th1/Th2 phenotype of the T cell immune response triggered by rBS*^CotB-HcG^*, we found that the anti-HcGAPDH IgG2a was 2.07 folds higher than the anti-HcGAPDH IgG1 (*p* < 0.005), indicating a Th1 dominated T cell immune response (Fig 4d). We also evaluated the levels of anti-HcGAPDH secretory IgA (sIgA) from intestinal epithelial cells and plasma cells, which could protect animals from pathogen infection by mucosal immunity. The results showed that anti-HcGAPDH sIgA was significantly induced in intestinal mucus of rBS*^CotB-HcG^* mice in comparison to that of Ctrl (*p* < 0.01) (S1 Fig). Genes representing Th1 activation (IFN-γ, IL-2, IL-12, and T-bet) and those of Th2 activation (IL-4, IL-6, IL-10, and GATA-3) in splenic lymphocytes were significantly induced by rBS*^CotB-HcG^* administration (Fig 4e), suggesting that rBS*^CotB-HcG^* stimulated mixed Th1/Th2 immune responses. Collectively, these data indicate that *B. subtilis* spores expressing the CotB-HcGAPDH fusion protein activated both humoral and cell-mediated immune responses in mice.

**Fig 4.**
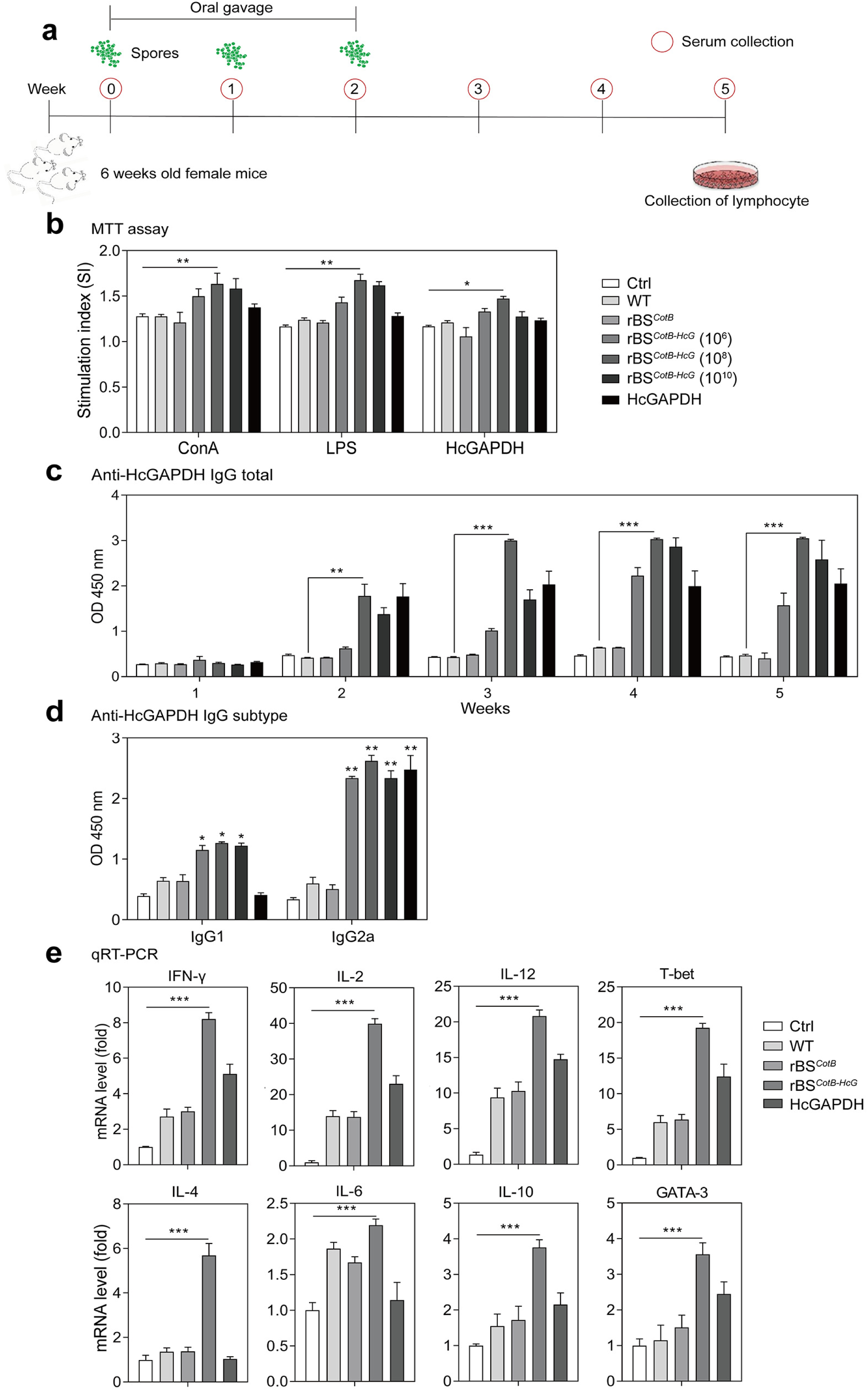
Recombinant *B. subtilis* spores expressing the CotB-HcGAPDH fusion protein induced both humoral and cell-mediated immune responses in mice. **a.** Schematic of the experimental protocol in mice. Six-week old female mice (n = 20 in each group) administrated with PBS (Ctrl), WT, rBS*^CotB^*, rBS*^CotB-HcG^* spores, or purified HcGAPDH protein at indicated dosages. Serum samples collected at indicated time points. **b.** Proliferation of splenic lymphocytes of mice (n = 6 in each group) measured by MTT assay. **c.** Anti-HcGAPDH IgG levels in sera of mice (n = 6 in each group) at different time points. **d.** IgG1 and IgG2a levels in murine sera at week 5 (n = 6 in each group). **e.** The mRNA levels of cytokine and transcription factor genes in splenic lymphocytes from mice (n = 6 in each group) measured by qRT-PCR. The dosage of rBS*^CotB-HcG^* was 1 × 10^10^ CFU. \**p* < 0.05, \*\**p* < 0.01, and \*\*\**p* < 0.005. All data are presented as means ± S. E. M. (standard error of the mean). Three technical replicates from a single experiment were used.

We next investigated the immune responses stimulated by the rBS*^CotB-HcG^* spores in sheep, one of the natural hosts of *H. contortus*. An in vivo experiment was carried out by gavage with PBS (control, Ctrl), *H. contortus* infection (Hc), wild-type (WT) strain or rBS*^CotB-HcG^* spores followed by *H. contortus* infection (Ctrl, Hc, Hc+WT and Hc+rBS*^CotB-HcG^*) (Fig 5a). Lymphocytes from the peripheral blood (PBLs) of sheep were isolated and cultured at day 7 after infection, a time point when *H. contortus* crawls to the abomasum and develops to the blood-sucking L4 stage. These cells were then stimulated with ConA, LPS or the purified HcGAPDH protein. Consistent with the murine results, proliferation of PBLs from sheep receiving Hc+rBS*^CotB-HcG^* in the presence of ConA or LPS was significantly higher than that from Hc group (p < 0.005) (Fig 5b). The purified HcGAPDH protein also stimulated significant proliferation of PBLs from these sheep in comparison to that from control sheep (*p* < 0.005). Administration of rBS*^CotB-HcG^* induced the anti-HcGAPDH IgG production (*p* < 0.005, compared with Ctrl) at week 2, and the level plateaued at week 4 and maintained until week 8 (Fig 5c). Meanwhile, Anti-HcGAPDH IgG was not detectable in the control sheep. Further, anti-HcGAPDH sIgA levels were significantly higher in intestinal mucus of Hc+rBS*^CotB-HcG^* sheep than that of the Hc sheep (*p* < 0.01) (S1 Fig). We also found that genes representing Th1 activation (IFN-γ, IL-2, IL-12, and TNF-α) and those of Th2 activation (IL-4 and TGF-β) in PBLs of Hc+rBS*^CotB-HcG^* sheep were highly activated (Fig 5d) even though expression of IL-6 and IL-10 did not change (*p* > 0.05). Collectively, these data show that rBS*^CotB-HcG^* stimulated strong humoral and cell-mediated immune responses in both mice and sheep.

**Fig 5.**
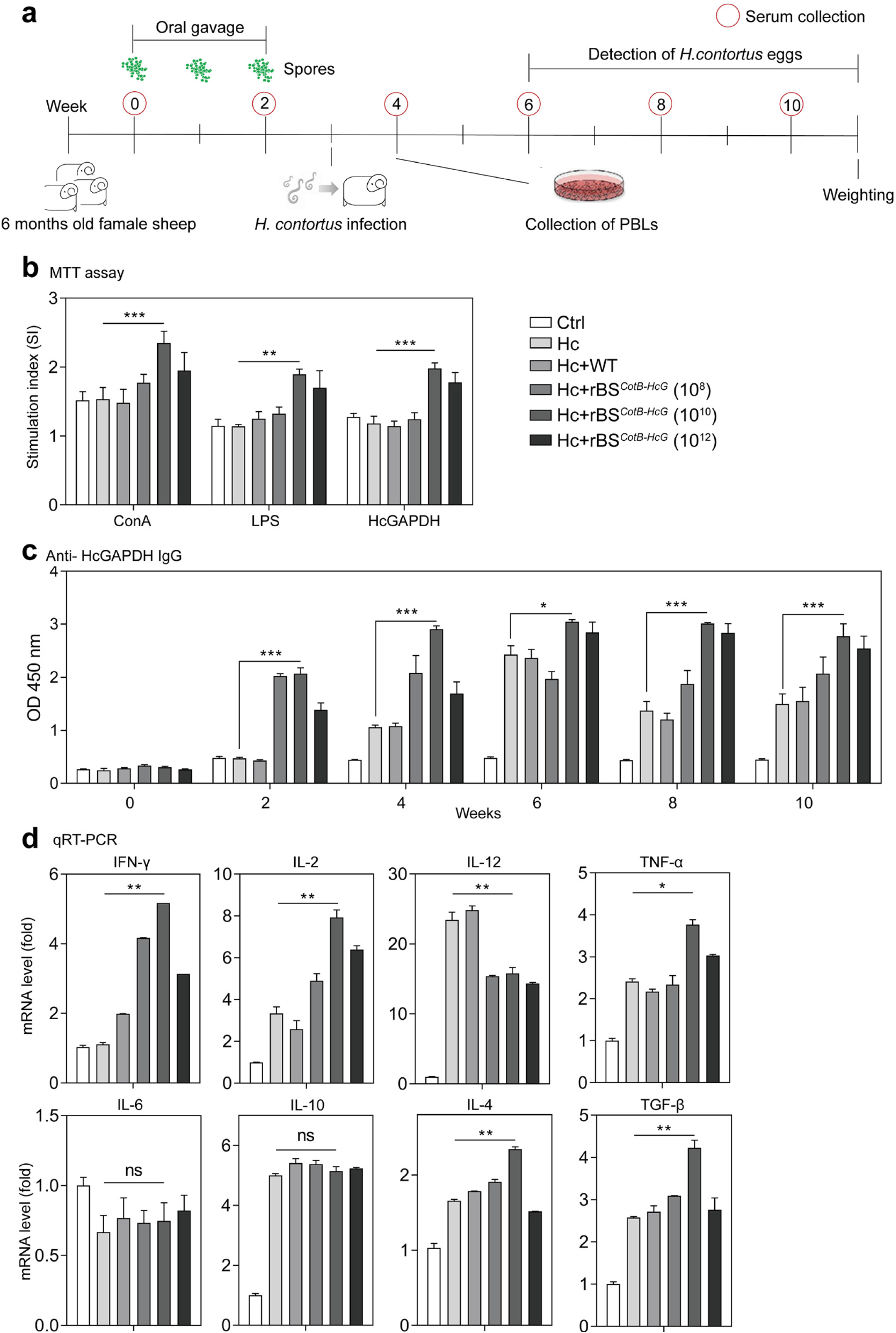
The CotB-HcGAPDH fusion protein expressing recombinant *B. subtilis* spores stimulated both humoral and cell-mediated immune responses in sheep. **a.** Schematic of the experimental protocol in sheep. Six-month old female sheep were fed by gavage with PBS (Ctrl), WT or rBS*^CotB-HcG^* spores at indicated dosages during the first 3 weeks, followed by *H. contortus* infection (n = 6 per group). Serum samples were collected at indicated time points. **b.** Proliferation of PBLs measured by MTT assay at week 4 (n = 6 in each group). **c**. Anti-HcGAPDH IgG levels in sera of sheep (n = 6 in each group). **d.** The mRNA levels of cytokine and transcription factor genes in peripheral blood lymphocytes (PBLs) of sheep (n = 6 in each group) measured by qRT-PCR at week 4. The dosage of rBS*^CotB-HcG^* was 1 × 10^12^ CFU. \**p* < 0.05, \*\**p* < 0.01, and \*\*\**p* < 0.005. All data are presented as means ± S. E. M. Three technical replicates from a single experiment were used.

### CotB-HcGAPDH recombinant *B. subtilis* spores promoted relative abundance of probiotic Bacilli in the abomasal microbiota in sheep

To investigate whether administration of rBS*^CotB-HcG^* affected abomasal microbiota of sheep in concomitant with *H. contortus* infection, 16S rRNA gene was sequenced from the abomasal mucus samples collected from the sheep of different treatment groups. Bacilli accounted for only less than 0.1% in the abomasal microbiota of sheep with *H. contortus* infection, compared with 4% of the controls (Fig 6a), consistent with our earlier findings (Fig. 1). Bacilli from Hc+rBS*^CotB-HcG^* sheep accounted for 3%, indicating that administration of rBS*^CotB-HcG^* could restore Bacilli depleted by *H. contortus* infection (Fig 6a). Community taxonomic system composition analysis of Firmicutes indicated that administration of rBS*^CotB-HcG^* increased the relative abundance of Lactobacillales (Fig 6b). Specifically, the Ctrl, Hc, Hc+WT and Hc+rBS*^CotB-HcG^* had a ratio of Lactobacillales abundance of 19.6%, 0.1%, 3.9% and 76.8%, respectively, in taxonomic composition of Firmicutes (Fig 6c). These results indicate that administration of rBS*^CotB-HcG^* spores improved the composition of the microbiota by increasing the ratio of probiotic species in the abomasum of sheep infected with *H. contortus*.

**Fig 6.**
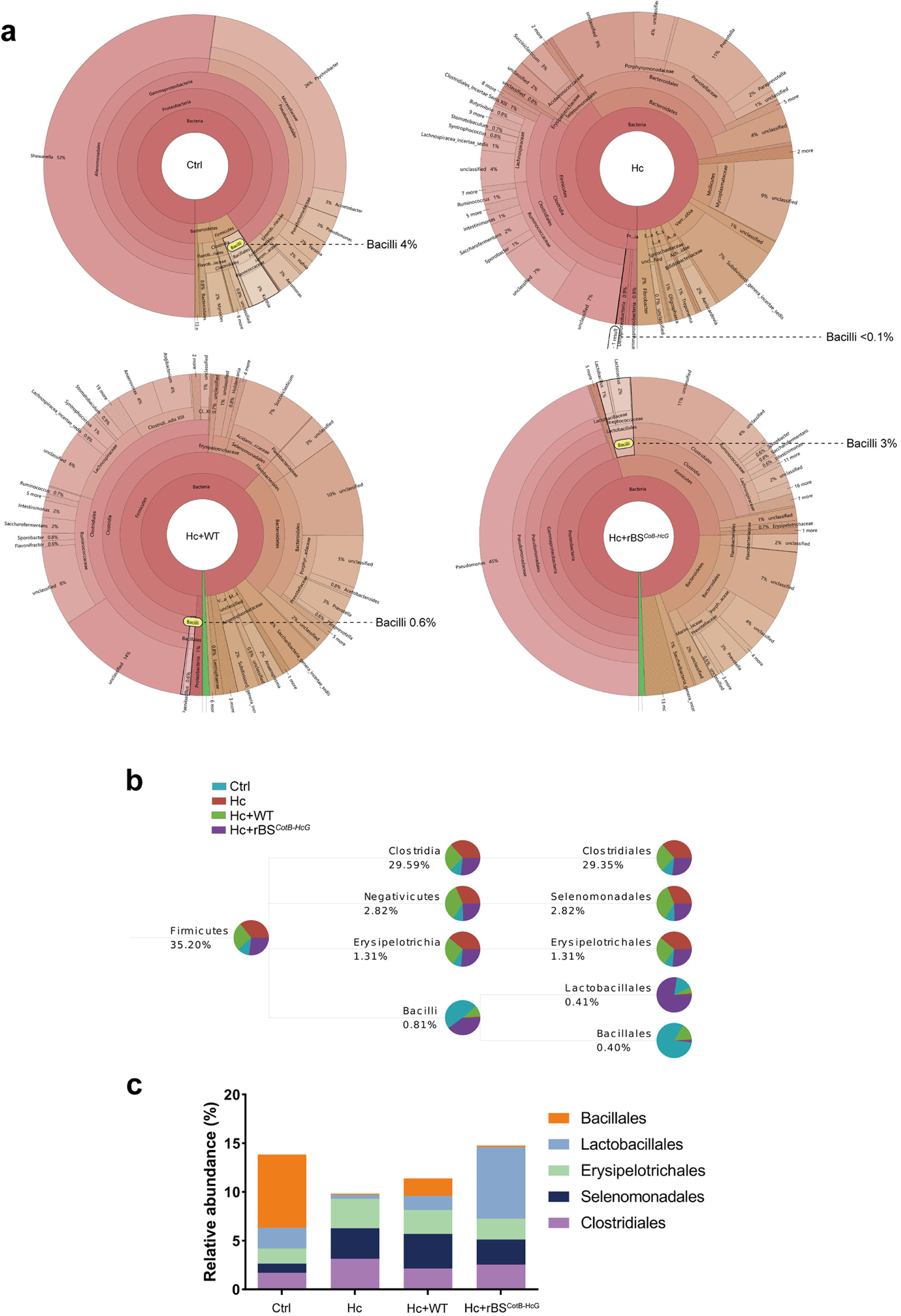
The CotB-HcGAPDH fusion protein expressing recombinant *B. subtilis* spores promoted relative abundance of probiotic Bacilli in the abomasal microbiota in sheep. **a.** Hierarchical analyses depicted in the form of a Krona plot. Six-month old female sheep orally gavaged with PBS (Ctrl), WT or rBS*^CotB-HcG^* spores in 10^12^ CFU/animal, followed by *H. contortus* infection (n = 3 in each group in all data). **b.** Heatmap of relative abundance at the class level of abomasal bacteria in sheep. **c.** Community taxonomic system composition analysis at the class level in sheep. **d.** Taxonomic composition of Firmicutes. The proportion of different color blocks indicates the relative abundance of different species.

### CotB-HcGAPDH recombinant *B. subtilis* spores protected sheep from *H. contortus* infection

To study the protective effect of rBS*^CotB-HcG^* on sheep against *H. contortus* infection, we measured the body weight and parasite loads of sheep. The average weight of the *H. contortus* infected sheep was only 49.5% of that of the non-infected control sheep, while sheep receiving rBS*^CotB-HcG^* at 10^10^ or 10^12^ CFU/animal followed by *H. contortus* infection could recover their body-weight back close to the controls. The wild-type *B. subtilis* spores also showed certain degree of protection against *H. contortus* infection, with 27.7% body weight gain compared to the infected sheep (Fig 7a). Next we determined parasite load by egg per gram feces (EPG) and adult worm counting. The sheep given 10^10^ CFU of rBS*^CotB-HcG^* /animal followed by *H. contortus* infection had their EPG dropped to 71.5% (Fig 7b). Their worm load also dropped to only 84.1% compared to the sheep infected with *H. contortus* (Fig 7c and Table 1). We also evaluated the infection by examining their abomasum. The surface of abomasum in the infected sheep was covered with worms and traces of parasite crawling. The numbers of worms and traces of parasite crawling in the Hc+rBS*^CotB-HcG^* sheep decreased compared with that of Hc group (Fig 7d). Further, the abomasum of infected sheep had intensive infiltration by mononuclear lymphocytes in mucosa, as compared with the un-infected sheep. No apparent infiltration was observed in Hc+rBS*^CotB-HcG^* sheep (Fig 7e). These data indicate that rBS*^CotB-HcG^* could offer effective protection of sheep from *H. contortus* infection by promoting immune responses and improving microbiota (Fig 7f).

**Fig 7.**
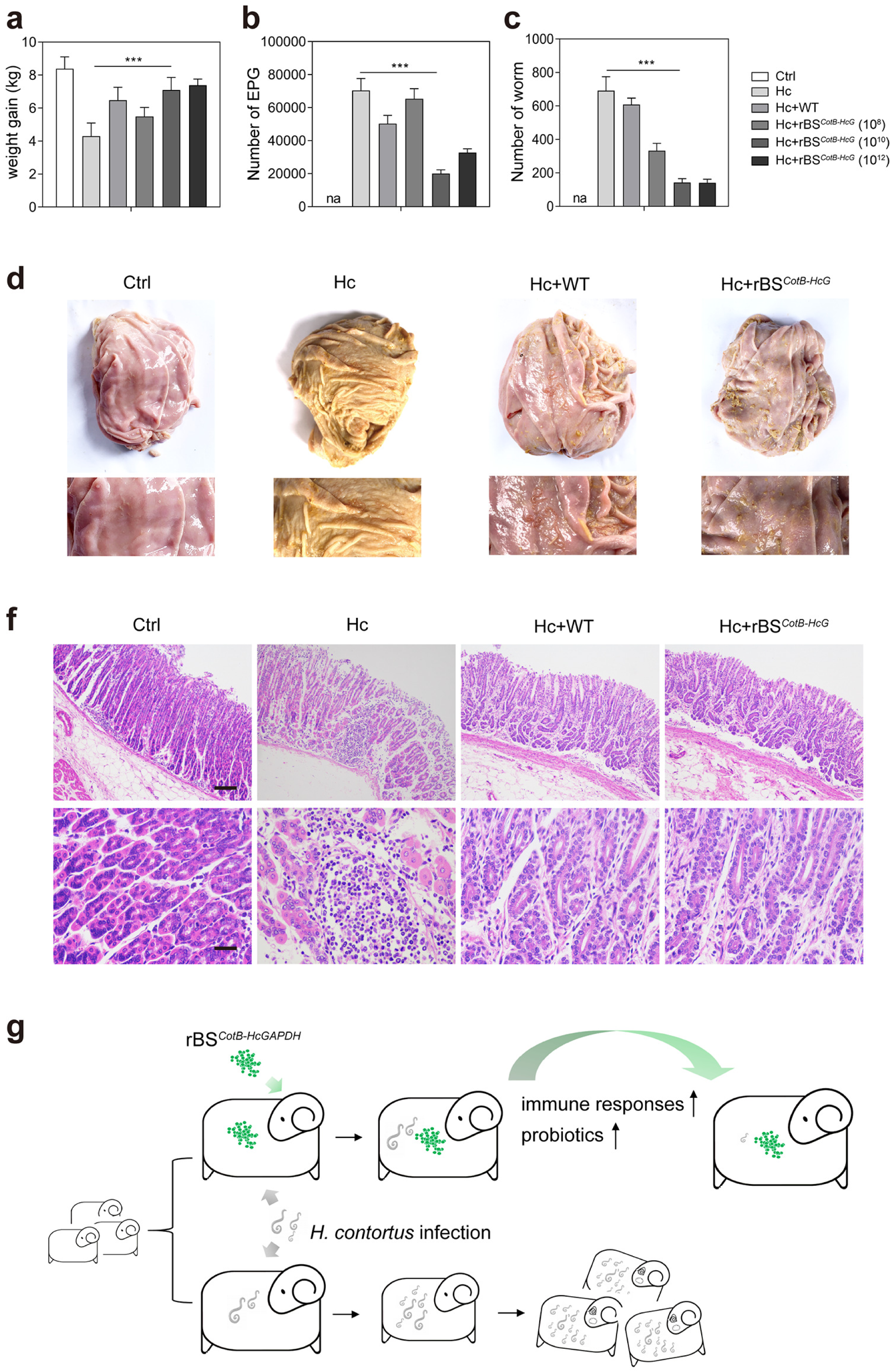
The CotB-HcGAPDH recombinant *B. subtilis* spores protected sheep from *H. contortus* infection. **a.** Average weight gain of sheep (n = 50 in each group). All data in this figure were collected from sheep followed the protocol shown in figure 5. **b.** Number of eggs per gram (EPG) of feces from sheep (n = 6 in each group). **c.** Number of worms from abomasum of sheep (n = 6 in each group). **d**. Representative pictures of abomasum in sheep (upper panel). Zoom-in images to show the *H. contortus* in abomasum in sheep (lower panel). **e.** HE staining of abomasum in sheep. All values in a, b, and c are presented as mean ± S. E. M. \**p* < 0.05, \*\**p* < 0.01, and \*\*\**p* < 0.005. **f.** Schematic of the protective effect of the CotB-HcGAPDH recombinant *B. subtilis* spores on *H. contortus* infection.

**Table 1.**
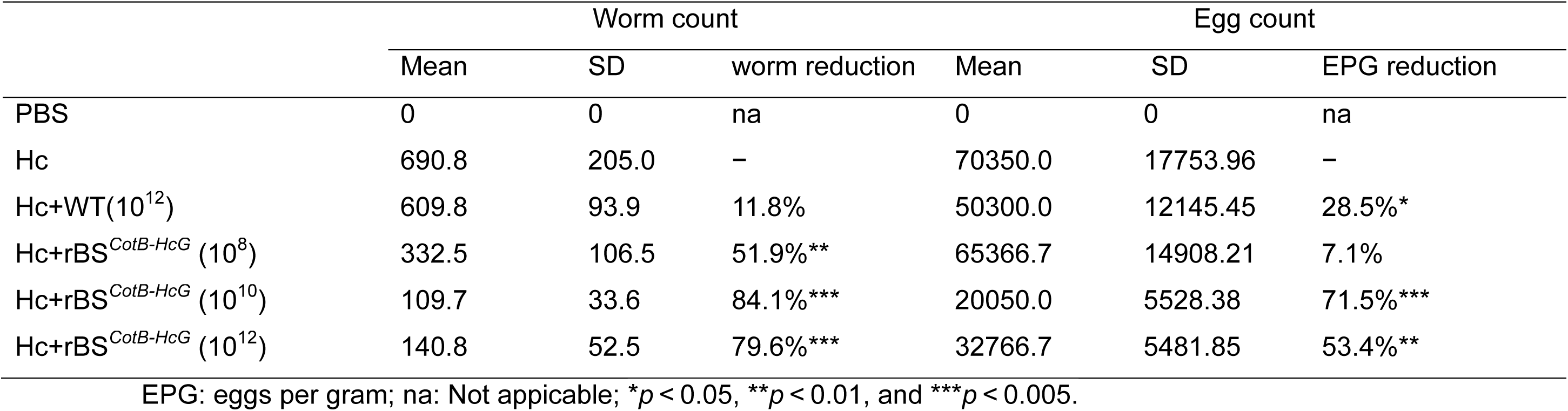
Worm reduction rate and EPG reduction rate in sheep with different treatments.

## Discussion

The goals of the current study were to evaluate protective capacity of HcGAPDH engineered on the *B. subtilis* spore surface in sheep against infection by *H. contortus* and to elucidate the immunologic mechanisms for its protection. A recombinant *B. subtilis* strain rBS*^CotB-HcG^* was developed by expression of the *H. contortus* protein HcGAPDH fused to CotB on the spore coat. Such recombination and heterologous expression did not change production and structure of the spores. However, the rBS*^CotB-HcG^* played an important role in regulating abomasal microbiota favoring the host sheep particularly when they were infected by *H. contortus* with perturbed microbiota in the abomasum. The rBS*^CotB-HcG^* induced Th1-dominated immune responses in a mouse model, a mechanism through which it can offer effective protection for sheep from *H. contortus* infection, and thus alleviated damages triggered by parasitic infections.

Oral vaccination has great potential for field use as large amounts of particulate materials can be delivered with low risk of adverse side effects [20]. More importantly, the probiotic-based strategy of vaccination could minimize the use of anthelmintics, thus reducing the risk of anthelmintic residues in food and minimizing the development of drug resistance in parasites. It is well known that the strategy of antigen delivery affects the levels of immune responses [21]. Dominant antibodies in the mucus are the first line of host defense against various pathogens invading the mucosa including parasites [22–24], by inhibiting the motility and adherence of pathogens in the mucus [25]. Also, sIgA and IFN-γ have a strong bactericidal effect in the early stage of infection [26]. However, whether oral administration is the best immunization strategy for *H. contortus* remains to be verified. Our data indicate that *B. subtilis* recombinant spores resisted the harsh conditions in the gastrointestinal tract and oral immunization with recombinant *B. subtilis* spores activated a strong mucosal immune response in the intestinal mucosa (S1 Fig).

Previous studies suggested delivery of the antigen via bacterial spores produced a Th1-biased cellular response, demonstrated by high levels of IgG2a [21]. Our results showed that murine splenic lymphocytes of rBS*^CotB-HcG^* -received mice expressed high levels of IL-2 and IFN-γ, which suggested these cytokines might contribute to non-protective Th responses against *H. contortus*. In addition, IL-12 is a key cytokine that induces Th1 type immune response [27]. The transcription factor T-bet is a major regulator of Th1 cell polarization [28]. Significant up-regulation of IL-12 and T-bet gene expression induced by rBS*^CotB-HcG^* indicated that *B. subtilis* spores mainly elicited Th1-type immune responses.

Interestingly, our results also indicate that the administration of rBS*^CotB-HcG^* in sheep induced the Th2 type immune responses besides Th1, as shown by up-regulation of cytokine IL-4 and TGF-β, implying that there may be mixed Th1/Th2 immune responses in sheep [29, 30]. A possible explanation is that such mixed immune responses are jointly activated by spores and HcGAPDH antigenic protein. Alternatively, rBS*^CotB-HcG^* by proteolytic cleavage might release soluble antigens including HcGAPDH, following their uptake by antigen-presenting cells (APCs), which may lead to presentation of a major histocompatibility complex class II-restricted manner (MHC-II) for generation of Th2 type immune responses [31]. IL-4 is a signal cytokine for the Th2 type response and is mainly responsible for IgE isotype switching [32]. The immunosuppressive cytokine IL-10 is responsible for inhibition of Th2 immune responses [33, 34]. In mice, we found a slight upregulation of cytokines (IL-4, IL-10) and Th2 type transcription factors (GATA-3), suggesting that rBS*^CotB-HcG^* could induce Th1/Th2 mixed immune responses. TGF-β is a functionally multidimensional cytokine that manipulates various immune activities differentially in various cell types and potentially regulates a wide range of biological processes. In sheep, the mRNA levels of TGF-β and IL-2 of lymphocytes in the peripheral blood were both increased. Some cytokines, particularly IFN-γ and TGF-β, have previously been proved to induce up-regulation of major histocompatibility complex class I-restricted manner (MHC-I) and MHC-II gene expression in different immune cells [35], which then stimulates production of antibodies and immune responses against parasitic pathogens [35]. Therefore, up-regulation of TGF-β gene expression in the sheep receiving of rBS*^CotB-HcG^* suggested that spores presenting HcGAPDH protein may activate the host immune responses against parasitic infections by stimulating the MHC-I and MHC-II antigen presenting pathways. Our results are consistent with those in an early study using different recombinant spores [36].

Here we have shown an example of the combination of a probiotic strain and a subunit vaccine that could enhance protection against *H. contortus* infection. *H. contortus* infection leads to significant decrease in the abundance of Bacillales in the abomasal microbiota. In addition, previous studies have shown that *B. subtilis* spores possess adjuvant property due to the co. However, a correlation of *H. contortus* infection with specific changes at the species or genus level of bacteria was not found. One possible explanation is that sequencing-based approach is set up to detect high-order taxonomic shifts rather than specific differences at the species or the genus level, consistent with an earlier report [7]. We further investigated probiotics at the level of Bacillales for controlling *H. contortus* infection. In recent years, *Bacillus* spp. are widely used as probiotics in the livestock industry, with some European Union-approved products available in the market. The most notable one is BioPlus2B from Christian Hansen [37]. The probiotic *B. subtilis* strain used as a carrier of passenger protein as vaccine has received attention because of its protective effects against various pathogens [20, 38–40]. Besides, previous studies have shown that *B. subtilis* spores possess adjuvant property due to the combination of antigens and spore surface [41, 42]. Many *Bacillus* strains are safe for sheep [43]. Therefore, they are likely suitable for use in sheep feeds. We have used these findings and attempted to establish a link between recombinant *B. subtilis* and prevention of *H. contortus* infection, by showing that the recombinant strain rBS*^CotB-HcG^* can offer significant protection against *H. contortus*.

Mechanistically, our working hypothesis (Fig 8) is that rBS*^CotB-HcG^* could activate T helper lymphocytes by APCs and stimulate up-regulation of IL-2 that might synergistically activate B lymphocytes to transform into plasma cells for generation of anti-HcGAPDH IgG antibodies. The spores could also stimulate the intestinal epithelial cells and plasma cells to produce anti-HcGAPDH sIgA, which would facilitate proliferation of eosinophils and up-regulation of TGF-β, resulting in parasite clearance. More importantly, the HcGAPDH is a key protein in inhibition of host complement activation. *H. contortus* living inside the host releases HcGAPDH that is involved in evasion of the host immune system. Thus, administration of rBS*^CotB-HcG^* could induce anti-HcGAPDH IgG and sIgA to block immune evasion of *H. contortus*.

**Fig 8.**
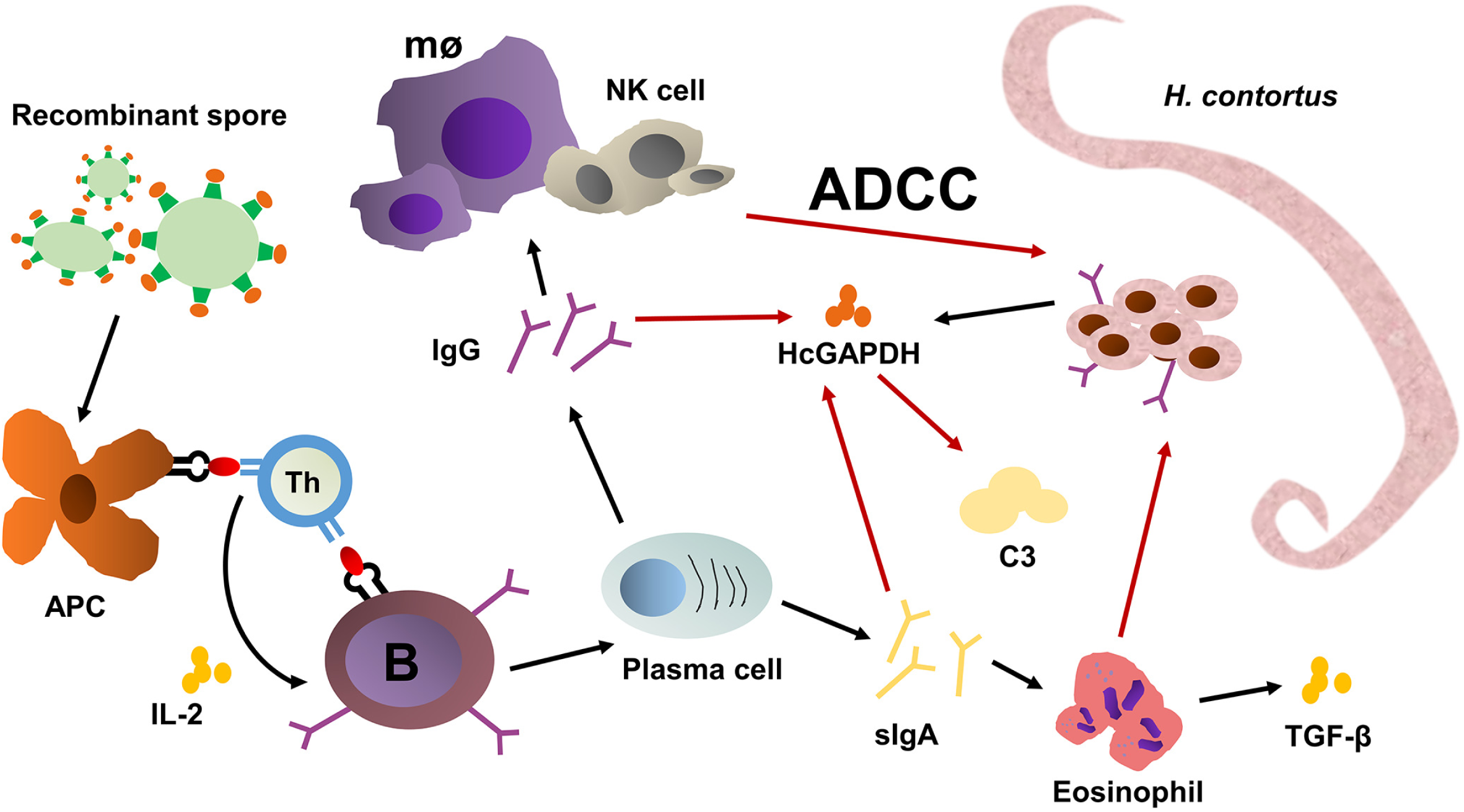
The hypothetic scheme of the recombinant spores expressing the CotB-HcGAPDH fusion protein in protecting sheep from *H. contortus* infection by ADCC effect. **a.** APC, antigen-presenting cells refer to a type of immune cells capable of ingesting, processing and processing antigens, and presenting the treated antigens to T and B lymphocytes. Th, helper T cell could help B cell produce antibodies. B, B lymphocyte. Mø, macrophages. ADCC, antibody-dependent cell-mediated cytotoxicity.

## Materials and Methods

### Ethics approval

Sheep (6 months of age; Huzhou, China) were maintained under helminth-free conditions in facilities at Zhejiang University. The procedures for animal maintenance and experiments were approved by Zhejiang University (permit no. ZJU20160239). All animal experiments were performed in accordance with guidelines for the care and use of laboratory animals and the experiments were approved by Zhejiang University Experimental Animals Ethics Committee.

### Parasite

*H. contortus* Zhejiang strain was kept at the Veterinary Parasitology Laboratory, Zhejiang University and maintained by serial passage in helminth-free sheep. Infective L3 larvae (iL3s) were obtained by incubation of eggs for 14days at 28 °C.

### 16S rRNA sequencing

Six-month old female sheep were orally infected with 5, 000 *H. contortus* iL3s, and euthanized at 14, 31 or 62 days post infection (DPI). The control sheep received 1 ml of PBS by oral gavage and euthanized at 62 DPI. These sheep were housed in separated areas to avoid cross-contamination within the university facility under the same environmental conditions. Ten ml of abomasum fluids was collected from each sheep by squeezing the whole abomasum within 20 min of euthanasia. The abomasal fluids were centrifuged at 5, 000 ×g for 5 min at 4 °C. The supernatants were re-centrifuged at 12,000 × g for 10 min at 4 °C and the pellet of each sample was used for 16S rRNA gene sequencing of abomasal microbiota by the Illumina MiSeq® platform (Sangon Biotech, China). Raw sequence files have been deposited in the Sequence Read Archive database under the project SRP217048. <u>Python v1.2.2</u> was used to analyze both heatmap and community taxonomic system composition. <u>LEfSe v1.1.0</u> was used to analyze taxonomic cladogram. <u>Krona v2.6.1</u> was used for hierarchical analyses.

### Plasmid construction

The 1,023 bp coding sequence (CDS) of HcGAPDH was amplified from the total cDNA of *H. contortus* by PCR using primers previously described [9]. PCR products were cloned into the pET-32a vector (Takara, China) via *Hind* III and *EcoR* I sites. The pET32a-HcGAPDH was sequenced (BioSune, China).

To generate a recombinant spore with a fusion protein of CotB-HcGAPDH, the genomic DNA of *B. subtilis* strain 168 was used as template to amplify the fragment of *CotB* gene (1,088 bp) containing the promoter sequence (263 bp) and *CotB* N-terminal partial CDS (825 bp) based on available sequences on NCBI (Reference Sequence: NC_000964.3) using the primers as listed in S1 Table. PCR products were cloned into the pMD 18-T vector (Takara, China) at *BamH* I and *Hind* III sites. The *CotB*-*HcGAPDH* was amplified by PCR with primers (S1 Table). The fused fragment of *CotB-HcGAPDH* was subcloned into *E. coli*-*B. subtilis* shuttle vector pDG364 (Miaolingbio, China) at *BamH* I and *EcoR* I sites, resulting in pDG364-CotB-HcGAPDH plasmid. The control plasmid pDG364-*CotB* was constructed by cloning *CotB* gene directly into pDG364 vector. All the plasmids were confirmed by sequencing (BioSune, China).

### Expression of recombinant proteins

The recombinant vector pET32a-HcGAPDH was transformed into *E. coli*. BL21 strain. The transformants were cultured at 37 °C until the OD600nm value reached approximately 0.6 and were then induced with 0.5 mM isopropyl β-D-1-thiogalactopyranoside (IPTG) at 37 °C. The pellets were collected by centrifugation at 8,000 ×g for 10 min, resuspended in buffer (0.01% digitonin, 10 mM Pipes, pH 6.8, 300 mM sucrose, 100 mM NaCl, 3 mM MgCl2, and 5 mM EDTA) with proteinase inhibitors, and processed by sonication. The soluble His-tagged protein was purified using a HisTrap column (GE Healthcare Life Sciences, USA). Purity of the eluted protein was checked by SDS-PAGE gel staining with Coomassie Blue. The anti-HcGAPDH rabbit polyclonal antibodies (rAb) were prepared according to the previous method [15]. The purified protein HcGAPDH and anti-HcGAPDH rAb were stored at −80 °C.

The pDG364-CotB-HcGAPDH plasmid was linearized by *Kpn* I and transformed into *B. subtilis* strain 168 by electroporation [44]. The fusion gene *CotB-HcGAPDH* substituted for amylase E (amyE) gene in the genome of *B. subtilis* by homologous recombination. *B. subtilis* spores were prepared in 4 L of DSM for sporulation of the recombinant *B. subtilis* strain rBS*^CotB-HcG^* or rBS*^CotB^* as previously described [21]. They were purified by treatment with 4 mg/ml lysozyme followed by washing under stringent conditions in 1 M NaCl and 1 M KCl with 1 mM PMSF. Spores were treated at 68 °C for 1 h in water to kill any residual sporangial cells. The final concentration of spores was set at 1 × 10^12^ CFU/ml in PBS pH 7.4. The spores were kept at −80 °C until use for animal experiments.

### SDS-PAGE and Western blotting

The transformed *B. subtilis* strain containing pDG364-CotB-HcGAPDH (rBS*^CotB-HcG^*) was cultured in LB medium with 25 μg/ml chloramphenicol at 37 °C. Bacterial sporulation was achieved by incubating in DSM according to the exhaustion method [21]. Spores were harvested and analyzed by SDS-PAGE to evaluate the presence of HcGAPDH protein. Moreover, spore coat proteins were extracted from spores at 48 h of bacterial incubation in DSM medium using SDS-DTT extraction buffer (0.5% SDS, 0.1 M DTT, 0.1 M NaCl) as previously described [44]. The extracted proteins were subjected to 12% SDS-PAGE and then transferred onto Polyvinylidene Fluoride (Sigma, Germany). The immobilized filter was blocked overnight at 4 °C in 5% skimmed milk in PBST (PBS with 0.05 % (v/v) Tween-20). Anti-HcGAPDH rAb (1:1000 in PBST) was used to probe the membrane by incubating for 2 h at RT after five washes in PBST. Finally, the probed filter was incubated with HRP-conjugated goat anti-rabbit IgG (1:5000 in PBST) and visualized by ECL (Beyotime biotechnology, China).

### Immunofluorescence and flow cytometry assay

Five ml of sporulation cultures at 24 h, 48 h or 72 h of incubation were harvested and processed as previously described [45]. Samples were blocked with 5% bovine serum albumin (BSA) for 2 h at 4 °C followed by incubation with anti-HcGAPDH rAb (1:2,000 in PBST) for 2 h at room temperature. Naive pre-immunized rabbit sera (1:2,000 in PBST) was used as negative control. Fluorescein isothiocyanate (FITC)-conjugated goat anti-rabbit IgG (Invitrogen, 1:500 in PBST) was used as the secondary antibody. Samples were observed and photographed under fluorescent microscope (Olympus BX51, Japan).

A total of 1 × 10^5^ purified spores were washed in PBS for 3 times and incubated with anti-HcGAPDH rAb (1:500 in PBST) at 37 °C for 2 h. Naive rabbit sera (1:500 in PBST) was used as negative control. After 3 washes in PBS, the spores were incubated with FITC-conjugated goat anti-rabbit IgG (1:500 in PBST, Invitrogen) at 37 °C for 1 h. Spores were finally resuspended in 1 ml of PBS following 3 washes, and at least 1 × 10^4^ spores were examined by FC500 MPL flow cytometer (Beckman Coulter, USA). Expression of the CotB-HcGAPDH fusion protein was analyzed using FlowJo software (Tree Star, USA).

### Analysis of the production and the structure of recombinant spores

The purified spores of the wild-type strain and the recombinant strain rBS*^CotB-HcG^* were collected, fixed in 3% glutaraldehyde overnight at 4 °C followed by dehydration in gradient ethanol of 50%, 70%, 90% and 100%. After subsequent critical point drying and sputter coating, they were processed and photographed under a scanning electron microscope SU-70 (Hitachi, Japan). For transmission electron microscopy, the spores were fixed in glutaraldehyde overnight at 4 °C followed by incubation in 4% osmium tetroxide for 2 h. Afterwards, they were dehydrated in gradient ethanol (50%, 70%, 90%, and 100%), embedded and the ultrathin sections were mounted on a 230 mesh copper mesh, stained with 1% uranyl acetate-lead citrate. The spores were observed and photographed under a transmission electron microscope H-9500 (Hitachi, Japan).

To investigate whether production of spores of the recombinant strain rBS*^CotB-HcG^* was different from that of the wild-type strain, both strains were inoculated 1 L of DSM medium, cultured at 37 °C with constant shaking at 140 r/min. The number of viable bacteria and spores were then quantified.

### Animal experiments

Six-week old female BALB/c mice were purchased from the Zhejiang Academy of Medical Science (Hangzhou, China), raised in a sterilized room, and fed with sterilized food and water. By oral gavage, the mice were administrated 100 μl PBS (Ctrl) per mouse, spores of wild-type strain at 1 × 10^10^ CFU (WT), rBS*^CotB^* at 1 × 10^10^ CFU (rBS*^CotB^*), and rBS*^CotB-HcG^* at 1 × 10^6^, 10^8^, or 10^10^ CFU per mouse (rBS*^CotB-HcG^*), respectively. The mice in Ctrl, WT and rBS*^CotB-HcG^* groups were administrated for three consecutive days, followed by two boosting, each for three consecutive days, at a one-week interval. Mice of the HcGAPDH group were subcutaneously immunized with 200 μg of purified HcGAPDH emulsified in the complete Freund’s adjuvant, followed by two boostings with 100 μg HcGAPDH emulsified in the incomplete Freund’s adjuvant at a one-week interval. All mice were euthanized at week 5 after the last immunization. Lymphocytes were isolated from spleens and cultured for extraction of total RNA.

Six-month old female sheep were purchased from the Miemieyang Animal Husbandry Co., Ltd. (Huzhou, China). All sheep were housed indoor and provided with hays and whole corns as food and water *ad libitum*. Each sheep was, by oral gavage, administrated with 1 ml PBS as control (Ctrl), spores of the wild-type strain (WT) at 1 × 10^12^ CFU per sheep (Hc+WT), and spores of rBS*^CotB-HcG^* at 1 × 10^8^, 10^10^ or 10^12^ CFU per sheep (Hc+rBS*^CotB-HcG^*), respectively. The sheep were challenged with 5,000 *H. contortus* iL3s one week after the oral gavage. Serum samples were collected from the jugular vein of each animal every two weeks. All sheep were sacrificed at two months post infection. PBLs were isolated at 5 DPI using a sheep peripheral blood lymphocyte separation kit (Sangon Biotech, China). Body weight gain of each sheep was recorded as the difference in body-weight (kg) between week 11 and week 0. EPG was assayed at 14 DPI according to the modified McMaster method [28]. The numbers of *H. contortus* adult worms from abomasum in sheep were counted after euthanasia at week 11.

### Lymphocyte proliferation assay

As described previously [46], murine PBLs were stimulated with LPS (5 μg/ml, Sigma, Germany), ConA (10 μg/ml, Sigma, Germany) or purified HcGAPDH protein (15 μg/ml). The cells were evaluated for proliferation by MTT Assay Kit (Sangon Biotech, China) according to manufacturer’s instructions. Experiments with sheep PBLs were performed the same way as the murine lymphocytes except for the concentration of LPS, ConA and purified HcGAPDH protein at 10 μg/ml, 15 μg/ml and 25 μg/ml, respectively.

### qRT-PCR assay

Total RNA was extracted from PBLs. The cDNA synthesized by qPCR RT Kit (TOYOBO, Japan) was subjected to quantitative real-time PCR (qPCR) to measure the mRNA levels of cytokines and transcription factors using a SYBR Green PCR Master Mix (Applied Biosystems, USA) on a StepOnePlus Real-Time PCR System (Applied Biosystems, USA). The primers specific for mouse or sheep TGF-β, IFN-γ, IL-2, IL-12, IL-4, IL-6, IL-10, T-bet, or GATA-3 gene were listed in S1 Table.

### Determination of antibodies by ELISA

Serum samples were collected from each mouse weekly after administration of the spores. The intestinal mucus samples were collected at week 5 according to a method previously described [31]. The levels of anti-HcGAPDH IgG, sIgA, IgG1 and IgG2a were measured by ELISA. Briefly, ELISA plates (Bethyl, USA) were coated with 1 μg purified HcGAPDH protein diluted in the coating buffer (0.05 M carbonate-bicarbonate, pH 9.6) followed by incubation in 5% skimmed milk for 18 h at room temperature. After three washes in PBST, the plates were then incubated at 37 °C for 2 h in serum or mucus in 1:400 dilution in PBST. Subsequently, HRP-conjugated goat anti-mouse IgG (1:5,000 dilutions, Abcam, UK), goat anti-mouse IgA (1:5,000 dilutions, Abcam, UK), goat anti-mouse IgG1 or IgG2a (1:1,000 dilutions, Abcam, UK) were employed as the secondary antibodies. After 1 h of incubation the plates were washed again and 100 μl substrate solution 3, 3′, 5, 5′-tetramethylbenzidine (TMB, BD biosciences, USA) was added. After 5 min of incubation in dark, the reaction was stopped by adding 50 μl 2 M H_2_SO_4_, and plates were read 3 times at 450 nm in the model microplate ELISA reader (BIO-RAD, Japan). Negative controls (coated with naive sera) were included on each plate. The results were expressed as OD450nm values. Similar to the mice serum protocol described above, anti-HcGAPDH IgG and sIgA in sheep samples were analyzed by ELISA as well. The secondary antibody was HRP-conjugated rabbit anti-sheep IgG and rabbit anti-sheep IgA (1:5,000 dilutions, Abcam, UK).

### HE staining

The abomasum dissected from sheep were thin-sectioned and subjected to HE staining [30]. The tissue sections were observed under an optical microscope (Zeiss, Germany).

### Analysis of abomasal microbiota from sheep

Relative abundance of abomasal microbiota in sheep of the Ctrl, Hc, Hc+WT, Hc+rBS*^CotB-HcG^* groups were analyzed by 16S rRNA gene sequencing. Sampling and sequencing process were consistent with the previous protocol [47].

### Statistical analysis

Results were presented as mean ± S. E. M. (standard error of the mean). Means of continuous variables were tested with two-tailed Student’s *t* test. *P* value of <0.05 was considered statistically significant.

### Data availability

All the data supporting the findings of this study are available within the article and its supplementary files.

## Acknowledgments

We would like to thank the Zhejiang University graduate students Wen Tang, Mingxiu Zhao, Hui Zhang, Mi Lin, Danru Bu, Mengjiao Li, Lulu Chen, Yifan Fang, Fei Wu, Lingyun Mou, and Haohan Zhuang. They were involved in the *H. contortus* challenge infection and sheep blood collection, mouse spleen lymphocyte separation and other assistance of experiments. We gratefully acknowledge Yang Wang of Nanjing University of Science and Technology for providing technical guidance for *bacillus subtilis* electro transformation. We would also like to thank staff of the Center for Animal Experiment of Zhejiang University and the Center for Electron Microscopy of Zhejiang University for animal care and help on electron microscopy, respectively.

## Author Contributions

Conceptualization: Aifang Du. Data curation: Yi Yang, Guiheng Zhang. Formal analysis: Yi Yang, Guiheng Zhang. Funding acquisition: Aifang Du, Yi Yang. Investigation: Yi Yang, Guiheng Zhang, Jie Wu, Danni Tong, Yimin Yang, Hengzhi Shi. Project administration: Aifang Du. Software: Yi Yang, Guiheng Zhang, Lenan Zhuang. Supervision: Aifang Du. Validation: Yi Yang, Guiheng Zhang, Aifang Du, Lenan Zhuang, Chaoqun Yao, Yimin Yang, Xueqiu Chen, Jianbin Wang. Visualization: Yi Yang, Guiheng Zhang. Writing – original draft: Yi Yang, Guiheng Zhang. Writing – review & editing: Aifang Du, Lenan Zhuang, Chaoqun Yao, Jianbin Wang. All authors read and approved the final version of the manuscript.

## Supporting information

**S1 Fig. The CotB-HcGAPDH fusion protein expressing recombinant *B. subtilis* spores induced both humoral and cell-mediated immune responses in sheep. a.** Anti-HcGAPDH sIgA levels in intestinal mucous samples of mice (n = 6 in each group). **b.** Anti-HcGAPDH sIgA levels in intestinal mucous samples of sheep (n = 6 in each group).

**S1 Table.**
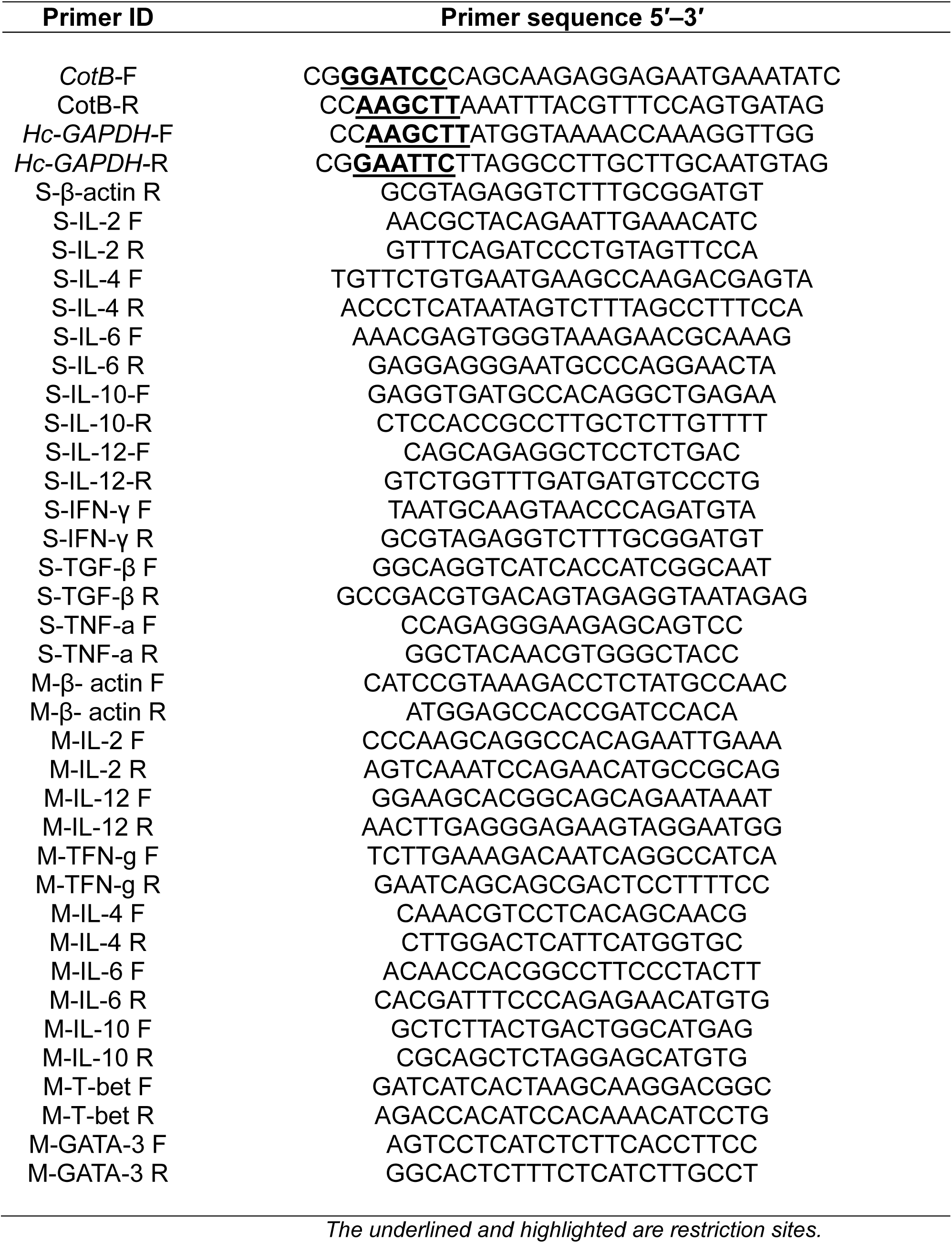
Primer sequence

